# Projected ecosystem responses to environmental changes associated with offshore wind farms and ocean warming

**DOI:** 10.64898/2026.08.26.747227

**Authors:** Bass Dye, Myron A. Peck, Johan van der Molen

**Affiliations:** Department of Coastal Systems, NIOZ Royal Netherlands Institute for Sea Research, Den Burg, 1790 AB, The Netherlands; Department of Animal Sciences, Marine Animal Ecology, Wageningen University & Research, Wageningen, 6700 AH, The Netherlands

## Abstract

Offshore wind farms (OWFs) are rapidly expanding to meet growing demands for renewable energy, with development expected to extend further offshore into deeper waters. Given this expanded human footprint, it is important to better understand the long-term ecological consequences of OWFs and how these may interact with ongoing climate change. We used the coupled hydrodynamic-ecosystem-biogeochemical water-column model (GOTM-ERSEM-BFM) to investigate ecosystem-wide responses to environmental changes associated with OWFs and climate warming. Specifically, we examined OWF-related scenarios of reduced benthic suspension-feeding activity, representing potential effects of contaminant emissions from OWFs, and reduced wind forcing, together with increased sea surface temperature. The scenarios were simulated individually and in combination to explore potential interactive effects. These scenarios were simulated at two contrasting locations in the North Sea, representing a well-mixed coastal site and a seasonally stratified offshore site. The coastal site exhibited comparatively modest ecosystem responses across the scenarios, whereas responses were generally stronger at the deeper offshore site. At the offshore site, changes in stratification altered vertical nutrient dynamics and contributed to pronounced differences in ecosystem responses between the surface and bottom layers. Our results demonstrate that ecosystem responses to OWF-related and climate-driven environmental changes are strongly dependent on local environmental conditions, suggesting that ecological consequences may differ substantially as wind farm development expands from shallower to deeper environments.

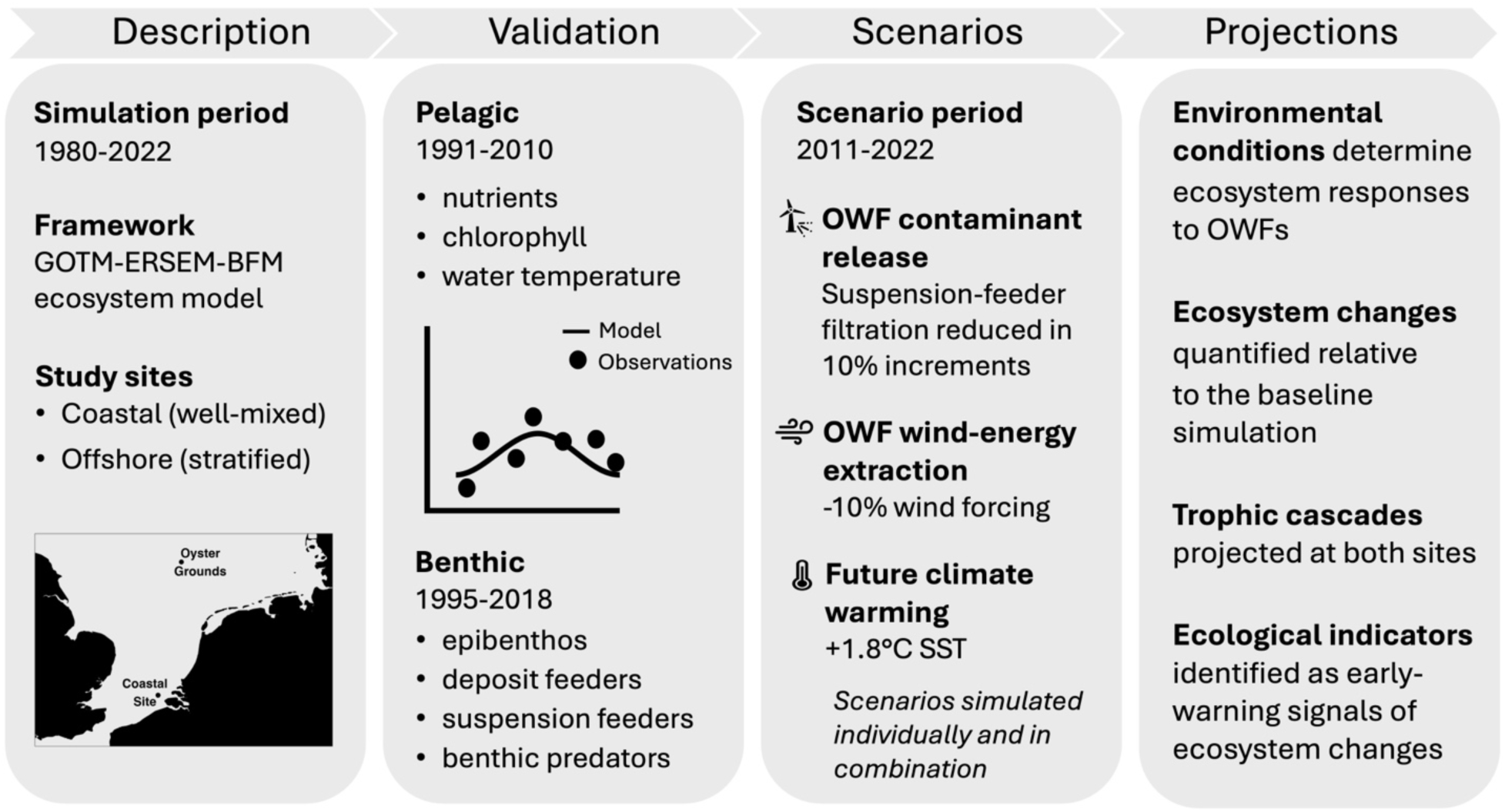

## 1 Introduction

Offshore wind energy is a rapidly expanding renewable energy sector, with global capacity expected to increase substantially in the coming years (Díaz and Soares, 2020). This expansion is expected to drive offshore wind farm (OWF) development further offshore into deeper coastal and shelf-sea environments (Renner, 2025). OWFs provide a major source of renewable energy, helping to reduce greenhouse gas emissions and mitigate climate-driven warming, which poses imminent threats to marine ecosystems (Ipcc, 2023; Hoegh-Guldberg and Bruno, 2010). Alongside their role in renewable energy production, OWF development can also modify marine ecosystems through multiple physical, chemical, and biological pathways. These include the introduction of artificial hard substrate (Degraer et al., 2020), alterations to local wind and hydrodynamic conditions through energy extraction (Christiansen et al., 2022; Akhtar et al., 2021; Christiansen and Hasager, 2005), and the release of contaminants from OWF infrastructure (Hengstmann et al., 2025; Czerner et al., 2025). In addition, broader climate-driven changes such as ocean warming (Kristiansen et al., 2024) may interact with OWF-associated environmental changes, with potentially important consequences for marine ecosystem dynamics.

Suspension feeding organisms play an important role in coastal and shelf ecosystems by coupling benthic and pelagic zones (Graf, 1992; Griffiths et al., 2017). Through filtration, suspension feeders remove particles from the water column and transfer organic matter to the seabed, ultimately influencing nutrient cycling, light penetration, and secondary production (Slavik et al., 2019; Mavraki et al., 2020; Maar et al., 2009). After construction, OWFs support the growth of suspension-feeder communities both on turbine foundations and in surrounding benthic habitats, representing multiple pathways through which changes in suspension-feeder filtration may influence ecosystem functioning (Degraer et al., 2020; Lefaible et al., 2019). A potential stressor associated with OWF development is the release of contaminants from corrosion protection systems (Wang et al., 2023), which are known to reduce filtration rates of suspension feeders (e.g., Mao et al., 2011; Kádár et al., 2001). These contaminants include a wide range of inorganic and organic compounds and heavy metals (Hengstmann et al., 2025; Czerner et al., 2025). Changes in filtration rates by suspension feeders can propagate through marine food webs and alter ecosystem functioning (De Borger et al., 2025).

In addition to potential changes in suspension-feeder filtration, OWFs can influence marine ecosystems by modifying the local physical conditions. By extracting energy from the atmosphere, OWFs reduce local wind forcing (Christiansen et al., 2022; Akhtar et al., 2021; Christiansen and Hasager, 2005). Reduced wind forcing can decrease vertical mixing and resuspension and strengthen water-column stratification, thereby affecting light conditions, nutrient availability, and ultimately primary production (Daewel et al., 2022; Van Der Molen et al., 2014). At the same time, ocean warming can strengthen thermal stratification while also increasing the metabolic rates of invertebrates, with consequences for nutrient cycling and trophic interactions (Doney et al., 2012; Thorpe et al., 2022). Importantly, these changes may interact with contaminant-induced changes in suspension-feeder filtration, potentially amplifying or dampening ecosystem responses relative to those arising from each pressure alone.

Predicting how OWF-associated and climate-driven environmental changes interact to influence marine ecosystems is essential for assessing the potential long-term ecological consequences of OWF expansion (Isaksson et al., 2025; Renner, 2025). Ecosystem models provide a valuable framework for representing key functional processes of ocean biogeochemistry and the lower trophic levels that structure marine food webs (Van Der Molen et al., 2013; Van Der Molen et al., 2014). Such models offer insight into the complex mechanisms governing marine systems, including trophic interactions and nutrient cycling dynamics (e.g., Griffiths et al., 2017), which are often difficult to resolve from observational and monitoring data alone (Skogen et al., 2024). Here, we use an ecosystem model to investigate the ecosystem-wide impacts of three scenarios and their interactions: (1) reduced benthic suspension-feeding activity, (2) reduced wind forcing, and (3) increased sea surface temperature. Specifically, we focus on the benthic suspension feeders to examine how changes in filtration rates propagate through marine ecosystems. The resulting changes in nutrient cycling and ecosystem structure were examined at two North Sea sites: a shallower, generally well-mixed coastal site with an existing OWF installation, and a deeper, seasonally stratified offshore site in an area under consideration for OWF development (Prinsen et al., 2022; Waldman et al., 2026) that has been extensively studied and modelled (e.g., Van Der Molen et al., 2013; Ruardij et al., 1997). We present simulated ecosystem responses and identify ecological indicators that may serve as early-warning signals of potential ecosystem shifts (Methratta, 2025).

## 2 Materials and methods

### 2.1 Study sites

Two study sites were selected to represent contrasting environmental conditions, ecosystem dynamics, and potential wind farm applications. The coastal site, located near the border of the Netherlands and Belgium, is an operational windfarm location (Fig. 1). This shallow, well-mixed location (22 m depth) represents a dynamic coastal environment characterized by strong tidal currents, high suspended sediment concentrations, and elevated primary productivity (Ivanov et al., 2020; Ivanov et al., 2021).

**Figure 1.**
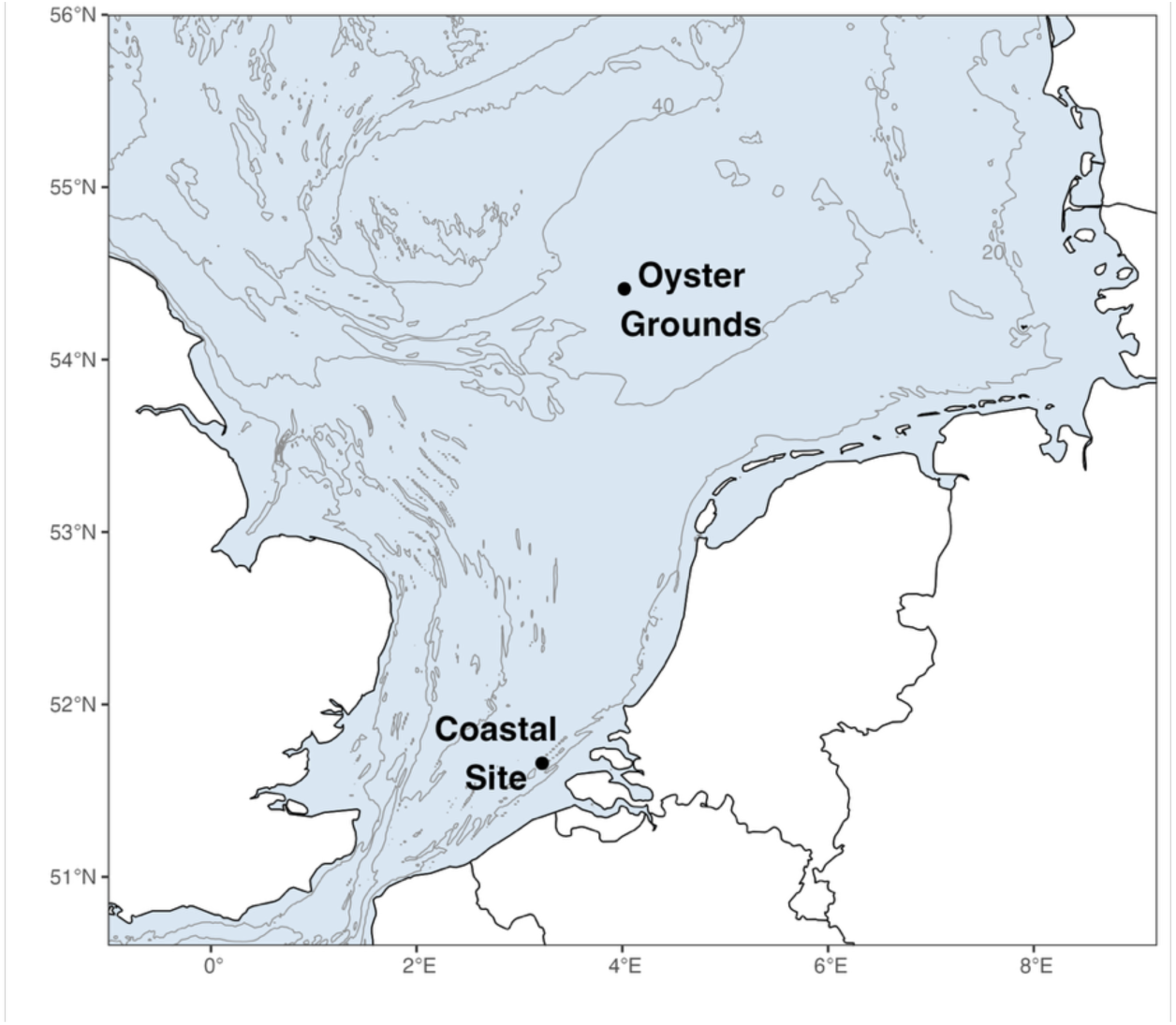
Map of the southern North Sea showing the locations of the coastal site and the offshore Oyster Grounds site used in the model simulations. Bathymetric contours indicate depth (m).

In contrast, the offshore Oyster Grounds site is deeper (45 m) and seasonally stratified, with weaker currents and lower suspended sediment concentrations (Van Leeuwen et al., 2015; Fig. 1). Wind farms are currently absent from the Oyster Grounds, although the area has been discussed as a potential site for future development (Prinsen et al., 2022; Waldman et al., 2026). Together, these locations represent contrasting hydrographic regimes of the southern North Sea, allowing assessment of potential ecosystem responses under both coastal and offshore conditions.

### 2.2 Observational data

Time-series observations were used to parameterize the model and evaluate its ability to reproduce observed ecosystem conditions. Monthly data from fixed-monitoring locations Walcheren 20 and Terschelling 135, part of the Rijkswaterstaat (RWS) Monitoring Waterstaatkundige Toestand des Lands (MWTL) program (https://waterinfo-extra.rws.nl/monitoring/), provided measurements of nutrients, chlorophyll, temperature, and salinity for the period 1991– 2010 at the coastal and offshore sites, respectively. These observations informed model parameterization and allowed assessment of model skill in reproducing seasonal dynamics across the measured variables.

Additionally, the model’s ability to reproduce observations of specific benthic functional groups was evaluated. However, these efforts were constrained by sparse sampling and limited proximity to the modelled sites. Additional annual data from the MWTL program for 1995-2018, collected from fixed-monitoring locations Walcheren 30 (10 km from the modelled coastal site) and Terschelling 100 (35 km from the modelled offshore site), provided observations of epibenthic predators, deposit feeders, suspension feeders, and infaunal predators for model comparison. While these observations enabled model assessment, the results should be interpreted cautiously given the temporal and spatial constraints.

### 2.3 Model description

The General Ocean Turbulence Model coupled with the European Regional Seas Ecosystem Model-Biogeochemical Flux Model (GOTM-ERSEM-BFM) was used to perform the simulations. GOTM-ERSEM-BFM is a one-dimensional mechanistic framework that couples water-column hydrodynamics with biogeochemical processes including nutrient cycling and food web dynamics. The model also incorporates a three-layer benthic module that simulates remineralization processes and a benthic food web composed of five functional groups.

#### 2.3.1 General Ocean Turbulence Model (GOTM)

The General Ocean Turbulence Model (GOTM; Burchard et al., 1999; http://www.gotm.net, last access: 4th February 2025) is an open source, one-dimensional water column model that resolves vertical hydrodynamic and thermodynamic processes. The model solves the one-dimensional Reynolds-averaged conservation equations for momentum, temperature, salinity, and turbulent quantities, allowing simulation of vertical mixing in the water column. Salinity was set to constant values of 33.5 and 35 PSU at the coastal and offshore site, respectively, representing average site conditions. The model was forced with meteorological hindcast data from the European Centre for Medium-Range Weather Forecasts (ECMWF; ERA-5) and depth-averaged tidal velocities reconstructed from harmonic analysis of an existing three-dimensional model (Van Der Molen et al., 2017), which provided external pressure-gradient forcing. Together with bed-shear stress, these forcings drive temperature, turbulence and currents in the simulations (Burchard et al., 2006; Karpouzoglou et al., 2020).

#### 2.3.2 European Regional Seas Ecosystem Model-Biogeochemical Flux Model (ERSEM-BFM)

The European Regional Seas Ecosystem Model-Biogeochemical Flux Model (ERSEM-BFM) used in the study is a development of the ERSEM III model (Baretta et al., 1995; Ruardij and Van Raaphorst, 1995; Ruardij et al., 1997; Ruardij et al., 2005; Vichi et al., 2007; Van Der Molen et al., 2017; last access: 1 November 2024). ERSEM-BFM is a coupled pelagic-benthic ecosystem model that describes biogeochemical fluxes within both the water column and the sediment, including the lower trophic levels of the marine food web. The model simulates the cycling of carbon, nitrogen, phosphorous, silicate, and oxygen, and allows for variable internal nutrient stoichiometry and chlorophyll content within functional groups depending on nutrient availability and physiological status. Within the one-dimensional model framework, nitrogen, phosphorous, and silicate are fully conserved. Nitrogen gas (N_2_) produced through denitrification is immediately returned to the system as nitrate via atmospheric deposition (Van Der Molen et al., 2013), while carbon and oxygen are exchanged through unlimited atmospheric pools at constant concentrations.

The model represents pelagic and benthic organisms using a functional group approach to resolve key ecological processes, trophic interactions, and nutrient cycling within the ecosystem. In the pelagic compartment, six phytoplankton groups are included (diatoms, flagellates, picophytoplankton, dinoflagellates, resuspended benthic diatoms, and *Phaeocystis* colonies) together with five zooplankton groups (suspension feeder larvae, omnivorous mesozooplankton, carnivorous mesozooplankton, microzooplankton, and heterotrophic nanoflagellates). The benthic compartment includes four macrofaunal (epibenthic predators, deposit feeders, suspension feeders, infaunal predators) and one meiofauna group (meiobenthos). Pelagic and benthic aerobic and anaerobic bacteria are also represented. The model also includes enhanced production of transparent exopolymer particles (TEP) by diatoms under nutrient stress, promoting the formation of macroaggregates composed of TEP, diatoms, and other suspended material. These aggregates increase particle sinking rates and enhance the supply of organic matter to the benthic system.

Suspended particulate matter (SPM) concentrations are simulated as a function of water and current driven resuspension processes. Significant wave height, period, and direction are calculated using a simple wave model based on the Sverdrup-Munk-Bretschneider method (Van Der Molen et al., 2014). Resuspension of detritus is coupled to the sediment resuspension, both of which influence the underwater light climate and consequently net primary production (Van Der Molen et al., 2017).

#### 2.3.3 Suspension feeder module within ERSEM-BFM

The suspension feeder functional type represents benthic suspension feeders that play an important role in coupling benthic and pelagic zones. Filtering activity is dynamically regulated by food availability and energetic profitability, occurring when the net energetic gain from food exceeds the cost of filtration. When food encounter rates are low relative to metabolic costs, activity is reduced, whereas high food availability promotes active filtration. Filtering is additionally limited by suspended sediment concentrations. This formulation allows the model to represent flexible feeding behavior rather than constant clearance rates. Non-digested particles are rejected as pseudofaeces and transferred to detritus pools. Mortality occurs from predation by the epibenthic predator functional group, a background and density-dependent component, as well as losses due to resuspension when bed shear stress exceeds a threshold. Reproduction can occur twice per year (primarily in spring but also in autumn) if favorable conditions are met, including active filtration, sufficient food availability, benthic temperatures exceeding 6°C, and low bed shear stress.

### 2.4 Model evaluation and scenario analysis

Simulations were conducted for the period 1980–2022, with the first 10 years treated as model spin-up. The remaining simulation period was used for model evaluation against observations and to assess ecosystem responses under the different scenarios.

#### 2.4.1 Model evaluation against observations

Climatological comparisons between surface-layer model simulations and pelagic observations were based on monthly averages over the 1991-2010 period. In contrast, model skill was evaluated by downsampling model outputs to the observation times. Specifically, daily averaged model outputs were compared with the corresponding dates of the monthly observations available for the period 1991–2010. An exception was made for chlorophyll, for which a ±7-day model window mean centered on each observation was used to better capture the simulated timing of the spring bloom. Several skill metrics were calculated to assess the model’s ability to reproduce the observed environmental conditions, recognizing that the relatively sparse, but long-term, monthly sampling of the pelagic observations may limit the achievable model skill.

Comparisons between modelled and observed benthic fauna were based on log-scale bias metrics, reflecting the spatial separation between observations and modelled sites (10-35 km) and the temporal sparseness of the annual benthic sampling. Over the 1995-2018 period, model bias was calculated as the mean log-ratio between modelled and observed values and subsequently back-transformed using the exponential function to obtain a bias factor. Consequently, the analysis focuses on regional-scale agreement in biomass magnitude rather than model skill, as assessed for the pelagic variables.

#### 2.4.2 Scenario analysis

Ecosystem changes resulting from the model scenarios (Sect. 2.5) were calculated over the final 12 years of the simulation (2011-2022). Chlorophyll and plankton dynamics were evaluated during the main productive period (1 March – 30 September), which captures the dominant spring and summer bloom dynamics, whereas the remaining variables were calculated over the full year. To assess changes in chlorophyll phenology, bloom timing and magnitude were evaluated separately for the spring bloom period (1 April – 30 June) and the subsequent late-season period (1 July – 31 December). The results include mean values calculated over the 12-year simulation period for the surface-layer pelagic model, unless otherwise specified (e.g., benthic layer), to facilitate comparison with observational data and to identify indicators that can be monitored.

### 2.5 Model scenarios

Three model scenarios (and their interactions) were simulated to discern ecosystem-wide impacts of OWFs under present and future environmental conditions and to assess the extent to which these scenarios influence ecosystem functioning.

#### 1) Pollutant release from OWFs

Corrosion protection systems used in OWFs, including protective coatings and sacrificial anodes, can release a wide range of inorganic and organic compounds into the marine environment (Kirchgeorg et al., 2018; Hengstmann et al., 2025; Ebeling et al., 2023; Ebeling et al., 2025; Reese et al., 2020). The magnitude and composition of these emissions are expected to vary over the operational lifetime of OWF turbines (Kirchgeorg et al., 2018). Exposure to such pollutants is known to reduce filtration activity in suspension feeders (e.g., Mao et al., 2011; Kádár et al., 2001). In the model, this effect was represented as a reduction in effective filtering capacity, implemented by scaling the volume filtered by suspension feeders. As a result, food encounter rates decrease, leading to lower feeding profitability and realized uptake (Sect. 2.3.3). Given the uncertainty in environmental exposure concentrations and associated toxicological responses (e.g., Alter et al., 2025; Ndugwa et al., 2026; Zonderman et al., 2025), we explored a range of impacts by reducing the filtering capacity in 10% increments to represent different, chronic levels of pollutant impact.

#### 2) Wind energy extraction by OWFs

OWF turbines extract kinetic energy from the atmosphere, reducing near-surface wind stress at the sea surface. Studies suggest that wind speeds are reduced by approximately 5-15% within OWFs (e.g., Christiansen et al., 2022; Akhtar et al., 2021; Christiansen and Hasager, 2005) which can influence primary production and other ecosystem responses (e.g., Daewel et al., 2022; Van Der Molen et al., 2014). We represent this effect by reducing wind forcing by 10%, implemented in the GOTM model using a wind scaling factor of 0.9.

#### 3) Future sea surface warming

Sea surface temperatures are projected to increase over the coming decades due to greenhouse-gas emissions (Kristiansen et al., 2024), with potential consequences for ecosystem and food web dynamics (Thorpe et al., 2022). Statistically downscaled CMIP6 projections suggest that mean sea surface temperatures at our study sites will increase by approximately 1.8 °C from present-day conditions (1995-2014) to mid-term conditions (2041-2060 mean) under the middle-of-the-road SSP2-RCP4.5 scenario (Kristiansen et al., 2024). We represent this effect by increasing the air temperature forcing to produce a mean annual sea surface temperature increase of 1.8 °C.

Each of the three scenarios was simulated individually and in combination (a total of seven scenarios) to explore potential interactive effects between pollutant exposure, wind energy extraction, and future warming.

## 3 Results

### 3.1 Model confirmation of pelagic observations

Overall, at the coastal site, the model reproduced the seasonal dynamics of temperature, nitrate, silicate, and chlorophyll, but performed less well for phosphate. The seasonality of temperature, nitrate, and silicate was captured in both timing and magnitude, particularly given the variability in the nutrient observations, including the wide range of spring nitrate concentrations (Fig. 2a, b, c). Chlorophyll dynamics were also reasonably represented, with the model reproducing the characteristic seasonal pattern (Fig. 2e). However, peak concentrations during the spring blooms were underestimated, and the modelled peak occurred slightly later than observed. This offset is small relative to the intra-monthly variability in the observations, particularly during the spring bloom period. In contrast, phosphate dynamics were not well reproduced. The model overestimates the observed background concentrations and fails to capture drawdown during depletion periods (Fig. 2d).

**Figure 2.**
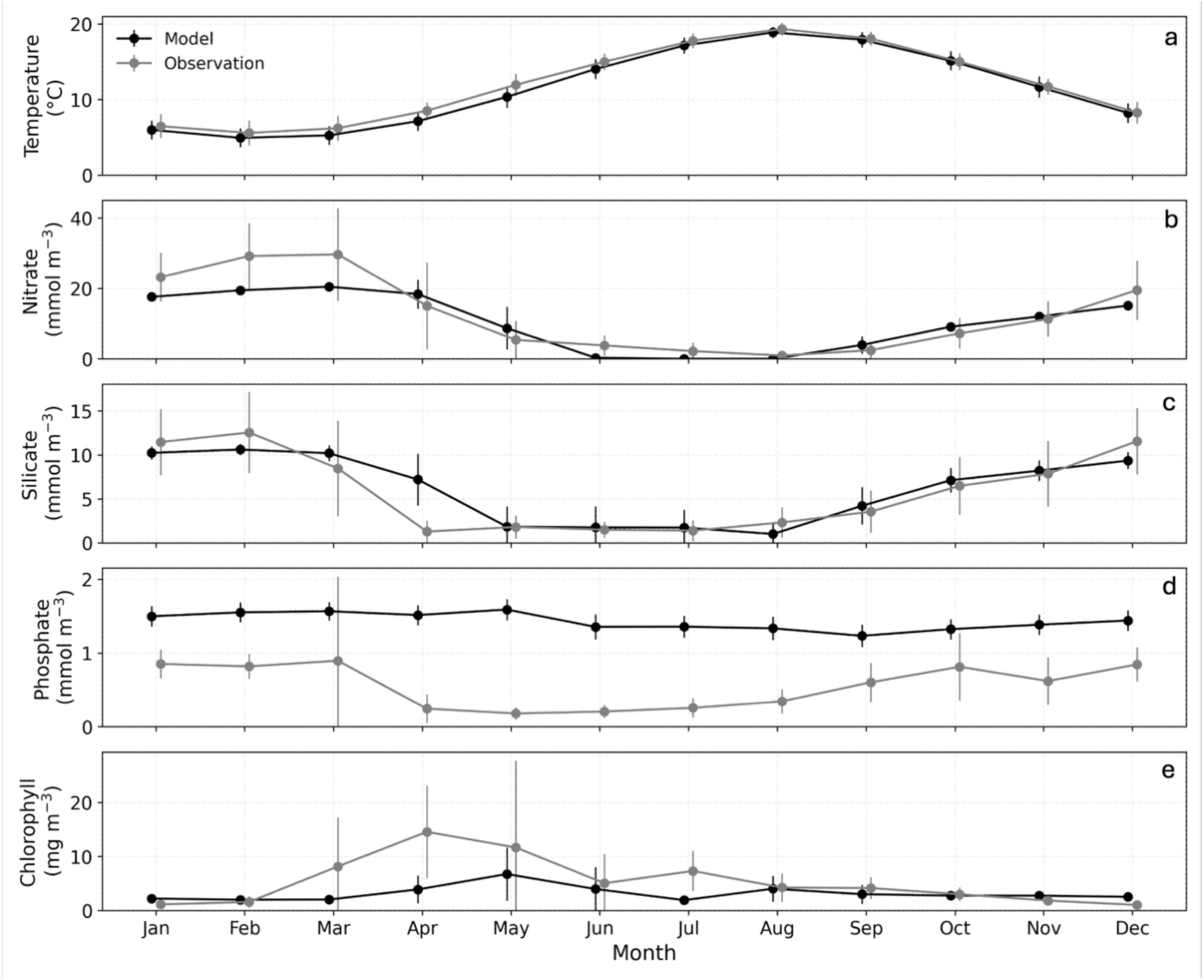
Comparison of observed and modelled climatological monthly means (± standard deviation) of temperature (a), nitrate (b), silicate (c), phosphate (d), and chlorophyll (e) at the coastal site for 1990–2010. Observations represent monthly measurements, while model values are monthly averages from the simulations over the same period. Observed and modelled means were slightly offset along the x-axis for visual clarity.

Model skill metrics at the coastal site are broadly consistent with the patterns observed in the climatological comparison (Table 1; Fig. 2). Temperature exhibits strong agreement with the observations (r = 0.95) while nitrate and silicate show relatively good agreement (r = 0.74 and 0.66, respectively). Additionally, each comparison yielded NRMSE values below 1, indicating that the model captures both the magnitude and variability of the observations (Table 1). Chlorophyll shows moderate agreement, with the model capturing general seasonal dynamics but with low correlation (r = 0.17) and a tendency to underestimate peak concentrations, consistent with the delayed and dampened spring bloom seen in the climatology (Fig. 2e). In contrast, phosphate is poorly represented, with near-zero correlation (r = -0.02) and a high NRMSE (2.09), reflecting discrepancies in both variability and magnitude.

**Table 1.**
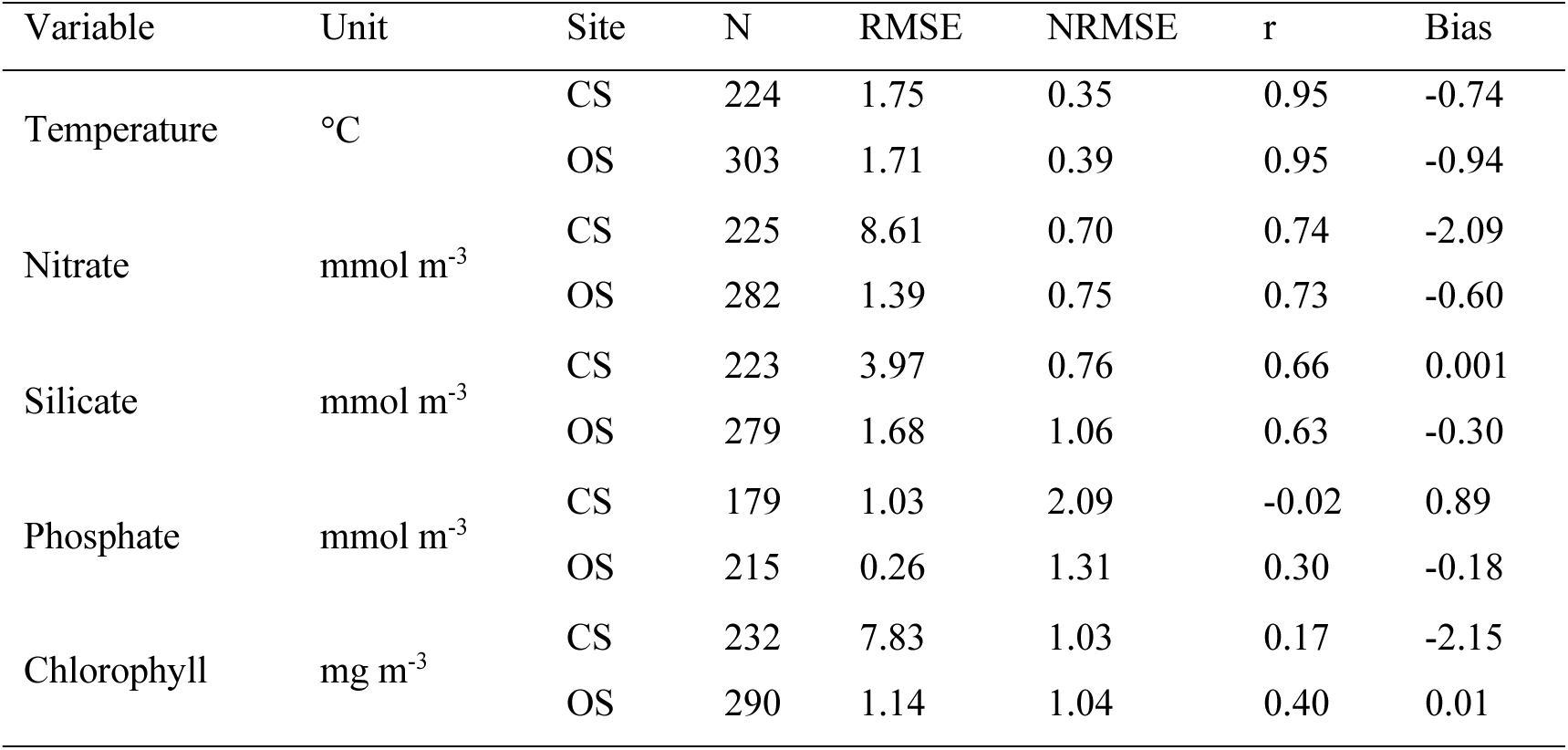
Model skill metrics for the coastal (CS) and offshore (OS) sites calculated from comparisons between model output and observations at matching sampling dates. N = number of observations; RMSE = root mean square error; NRMSE = normalized RMSE, RMSE divided by the standard deviation of the observations; r = correlation coefficient; Bias = mean difference (model – observation).

| Variable | Unit | Site | N | RMSE | NRMSE | $r$ | Bias |
| --- | --- | --- | --- | --- | --- | --- | --- |
| Temperature | °C | CS | 224 | 1.75 | 0.35 | 0.95 | -0.74 |
|  |  | OS | 303 | 1.71 | 0.39 | 0.95 | -0.94 |
| Nitrate | mmol m<sup>-3</sup> | CS | 225 | 8.61 | 0.70 | 0.74 | -2.09 |
|  |  | OS | 282 | 1.39 | 0.75 | 0.73 | -0.60 |
| Silicate | mmol m<sup>-3</sup> | CS | 223 | 3.97 | 0.76 | 0.66 | 0.001 |
|  |  | OS | 279 | 1.68 | 1.06 | 0.63 | -0.30 |
| Phosphate | mmol m<sup>-3</sup> | CS | 179 | 1.03 | 2.09 | -0.02 | 0.89 |
|  |  | OS | 215 | 0.26 | 1.31 | 0.30 | -0.18 |
| Chlorophyll | mg m<sup>-3</sup> | CS | 232 | 7.83 | 1.03 | 0.17 | -2.15 |
|  |  | OS | 290 | 1.14 | 1.04 | 0.40 | 0.01 |

At the offshore Oyster Grounds site, the model successfully reproduced the seasonal dynamics of the observations. Both the timing and magnitude of temperature and nutrient trends were generally well represented. Temperature dynamics were accurately captured (r = 0.95, Table 1; Fig. 3a). The model closely matched nitrate concentrations during the last three quarters of the year, though it underpredicted values in the spring (Fig. 3b); however overall agreement remained (r = 0.70, Table 1). Silicate was overpredicted in spring and underpredicted in fall (Fig. 3c), yet the model still captured the seasonal trend (r = 0.66, Table 1). The model also reproduced chlorophyll’s characteristic seasonal pattern both in timing and magnitude (Fig. 3e; r = 0.40, Table 1). Phosphate dynamics showed lower skill than the other modelled variables. While the model captured the general seasonal pattern of phosphate (Fig. 3d; r = 0.30, Table 1), the projections exhibited a bias as modelled concentrations dropped to near zero in summer, suggesting excessive limitation compared to the observations.

**Figure 3.**
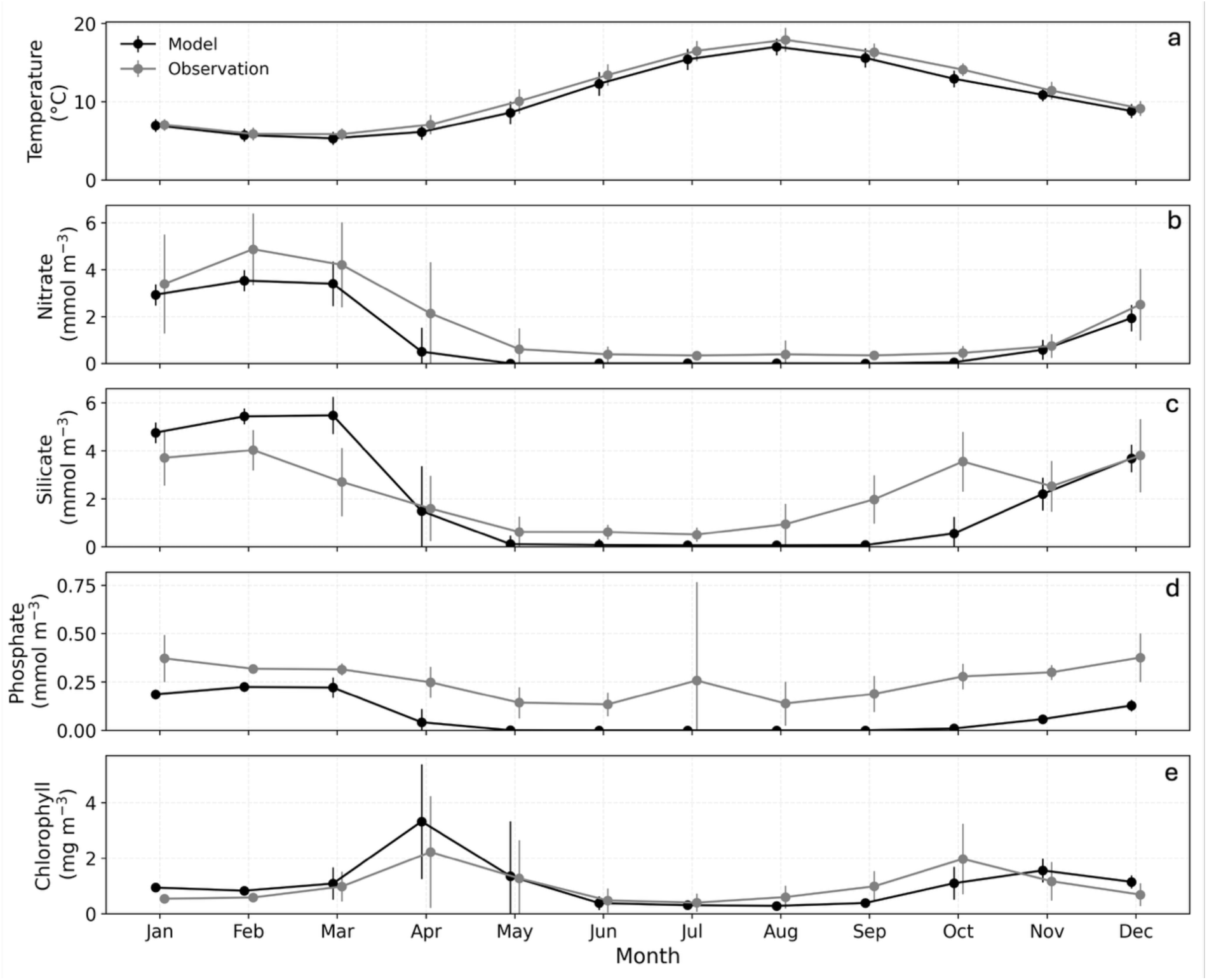
Comparison of observed and modelled climatological monthly means (± standard deviation) of temperature (a), nitrate (b), silicate (c), phosphate (d), and chlorophyll (e) at the offshore site for 1990–2010. Observations represent monthly measurements, while model values are monthly averages from the simulations over the same period. Observed and modelled means were slightly offset along the x-axis for visual clarity.

The model’s weaker representation of phosphate at both sites likely reflects the tight coupling of this nutrient with multiple interacting processes, including phytoplankton uptake, recycling, and benthic-pelagic exchange, that were difficult to constrain at the dynamic coastal site, although less challenging at the offshore site. Attempts to reduce phosphate concentrations at the coastal site to better match observations resulted in unrealistic system behavior, indicating that the chosen parameterization provides the best overall agreement with observed nutrient dynamics and ecosystem structure. Despite these discrepancies, the model captures the seasonal patterns of primary production and nutrient cycling at both sites.

### 3.2 Model confirmation of benthic observations

The model systematically overestimated benthic functional group biomass at both coastal and offshore sites, though the magnitudes of overestimation differed. Modelled biomass at the coastal site generally exceeded observations on average by 4 to 7 times across benthic functional groups, although deposit feeders showed the largest discrepancy at nearly 28 times (Table 2). Offshore estimates showed a smaller overall bias (∼2 times), though infaunal predators still exhibited elevated error (>5 times; Table 2). These discrepancies indicate differences in model to observation agreement across the habitat types and functional groups.

**Table 2.** Model bias for observed benthic functional groups at the coastal (CS) and offshore (OS) sites. Bias was calculated from comparisons between model output and observations at matching sampling dates. Observational stations were located approximately 10 km (CS) and 35 km (OS) from corresponding model locations. N = number of observations; Bias factor = exponent of mean log-ratio (model / observation).

| Variable | Unit | Site | N | Bias factor |
| --- | --- | --- | --- | --- |
| Epibenthic predators | mg C m<sup>-2</sup> | CS | 19 | 7.07 |
|  |  | OS | 19 | 1.82 |
| Deposit feeders | mg C m<sup>-2</sup> | CS | 19 | 27.88 |
|  |  | OS | 19 | 1.59 |
| Suspension feeders | mg C m<sup>-2</sup> | CS | 19 | 4.30 |
|  |  | OS | 19 | 2.32 |
| Infaunal predators | mg C m<sup>-2</sup> | CS | 19 | 7.16 |
|  |  | OS | 19 | 5.40 |

While the discrepancies identified throughout Sects. 3.1 and 3.2 reflect constraints inherent to both the model and observational data, the model nevertheless provides a suitable framework for exploring ecosystem responses to the scenarios considered (Sect. 3.3).

### 3.3 Individual scenario responses

All scenarios resulted in changes to ecosystem structure and functioning. Mechanistic interpretations are presented and supported by changes in key fluxes (e.g., grazing).

#### 3.3.1 Reduced filtration at the coastal site

At the coastal windfarm site, the simulations suggest that reductions in suspension-feeding filtering capacity can propagate through the pelagic food web via trophic cascades (Fig. 4a). Suspension-feeders remained viable even as filtering capacity was reduced to 30% of baseline, though biomass and reproductive output (larval stage) progressively declined throughout this range (up to -40%). This was accompanied by a decline in carnivorous mesozooplankton (up to -29%), coinciding with reduced availability of suspension-feeder larvae (up to -38%), a key prey item. The decline in carnivorous mesozooplankton weakened predation pressure on omnivorous mesozooplankton (up to -40%), allowing their biomass to increase (up to +96%). Flux analyses confirm that the increase in omnivorous mesozooplankton resulted in stronger grazing on diatoms (up to +64%), which contributed to a decline in diatom biomass despite the simultaneous reduction in direct grazing and filtering by the suspension feeders. As diatom biomass declined, silicate uptake decreased, leading to an increase in pelagic silicate concentrations (up to +14%). Reduced benthic aerobic and anaerobic silica dissolution further suggests reduced sedimentary silica recycling (up to -14%), consistent with lower diatom production and deposition. The other phytoplankton groups showed mixed responses, and total chlorophyll increased (up to +13%), indicating a shift in phytoplankton community composition rather than a strong change in total phytoplankton biomass. Consistent with this, net primary production showed little change (<4%). Despite changes in zooplankton community composition, total secondary production remained relatively stable (<5% change). Overall reduced filtration induced trophic cascades, restructuring zooplankton communities and indirectly altering phytoplankton and nutrient dynamics, while total ecosystem production remained relatively stable (Table 3).

**Figure 4.**
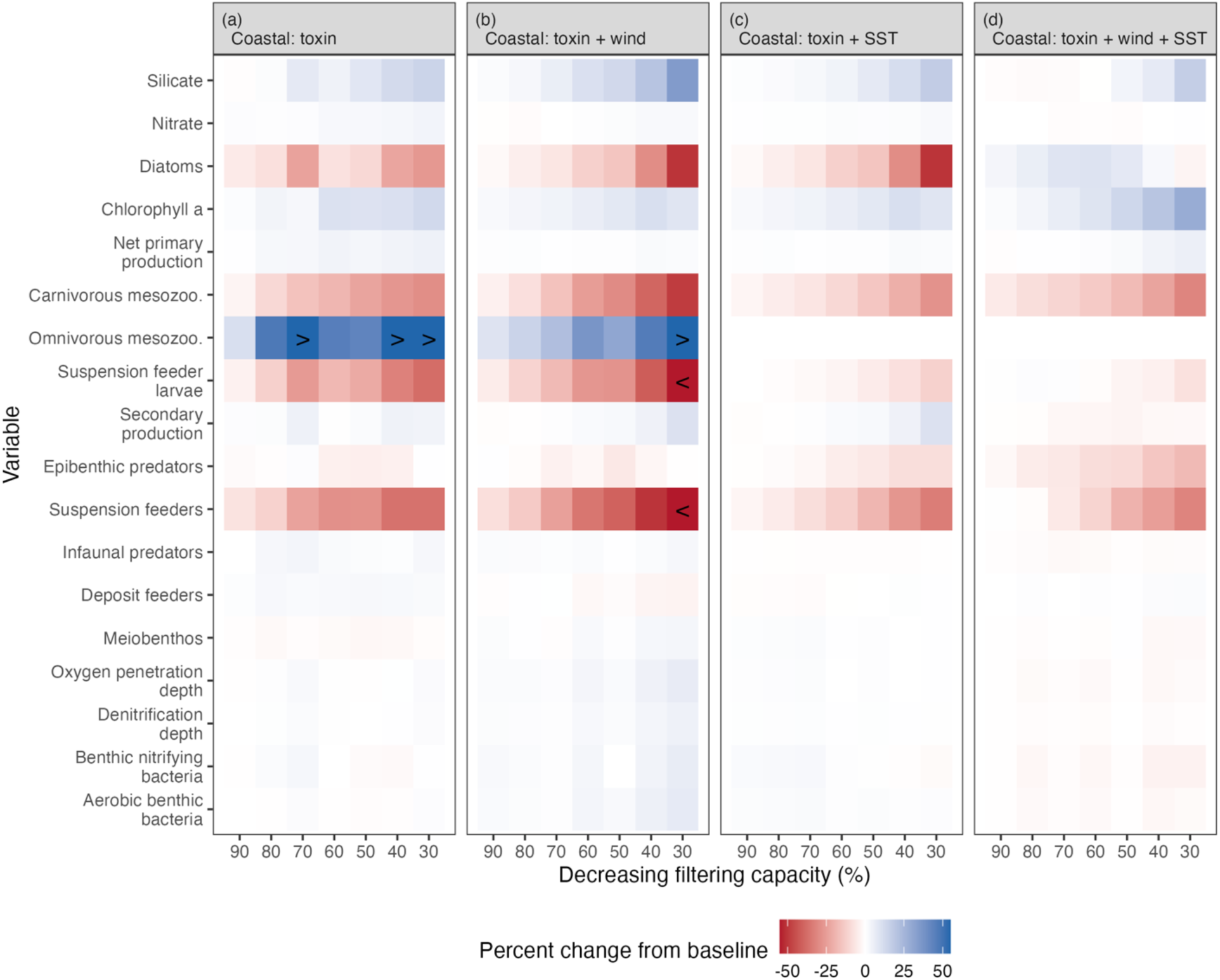
Simulated ecosystem responses to reduced filtering capacity across scenarios and their interactions at the coastal site, expressed as percent change relative to each scenario-specific baseline simulation with full filtering capacity. Pelagic variables are from the surface layer. Symbols (< and >) represent percentage changes below -50% and above +50%, respectively. Columns represent decreasing filtering capacity (from left to right) and rows represent ecosystem variables. Variable units are provided in Table 3.

**Table 3.**
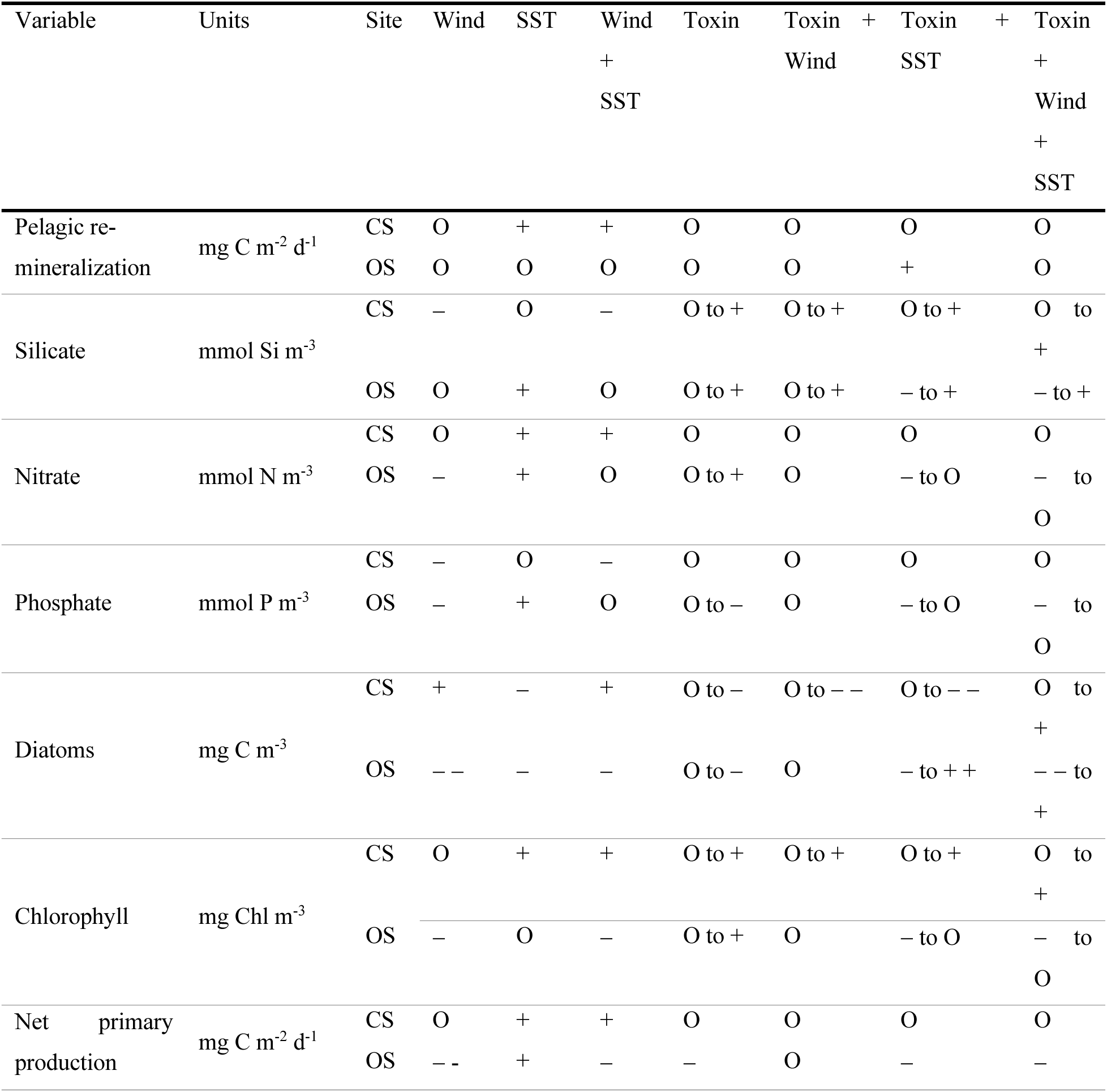
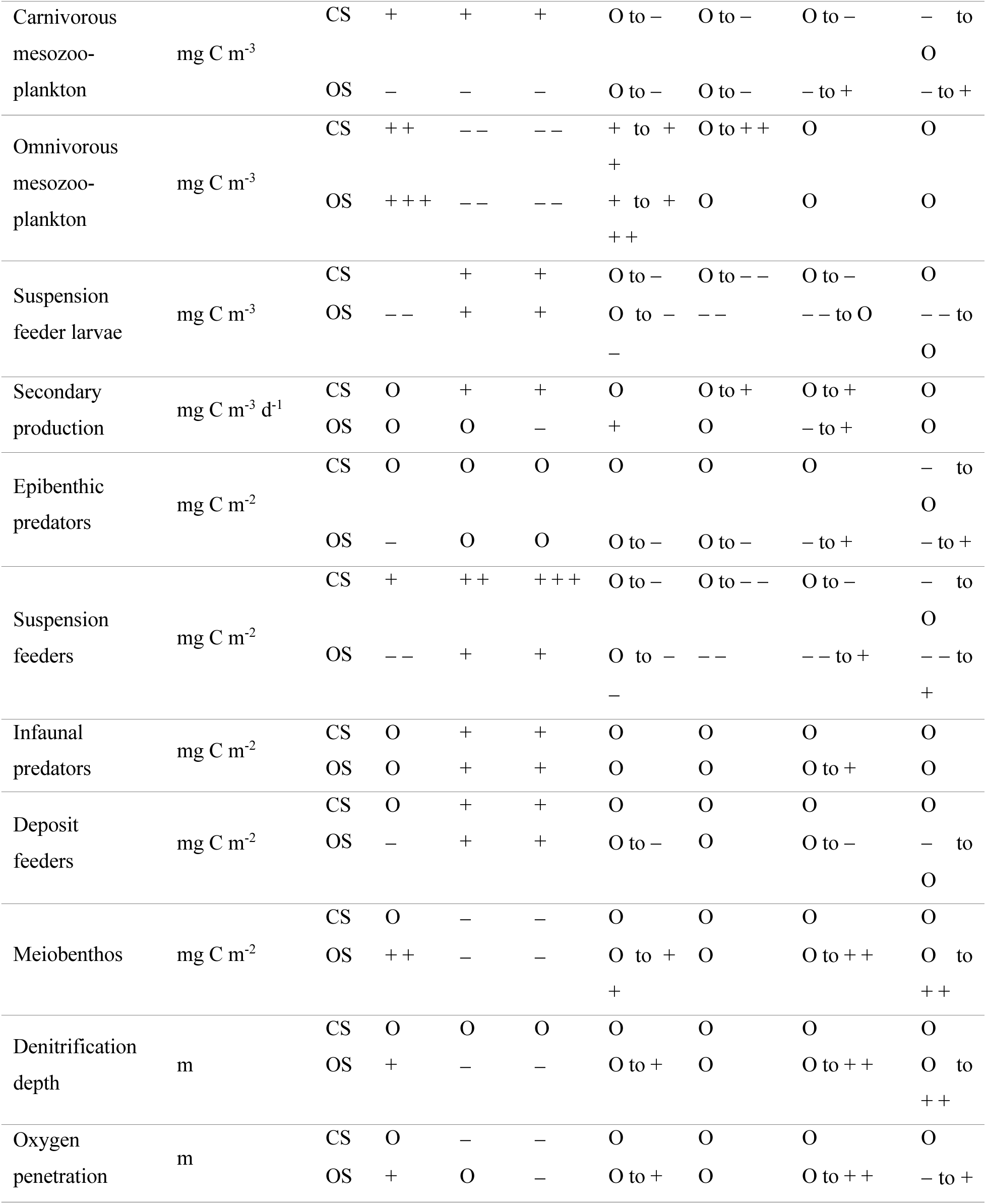
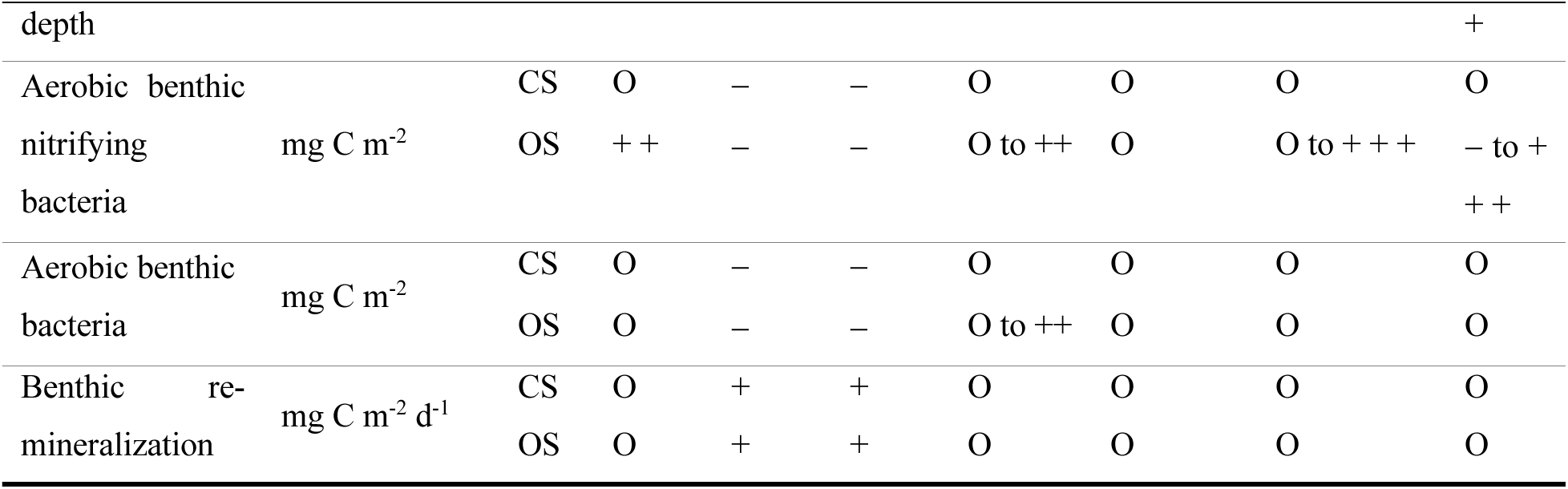
Comparison of scenario-driven changes in ecosystem state variables at the coastal (CS) and offshore (OS) sites. Pelagic variables are from the surface layer. The number of (+) and (–) symbols indicates the direction and relative magnitude of percentage changes (+/– = 10-49.9%; + +/– – = 50-100%; + + +/– – – = >100%), while changes of less than ±10% are denoted with (O). Changes are relative to the baseline scenario, except for the Toxin + scenarios, which are relative to the specific scenarios(s) without any filtering limitation. For the toxin scenarios, symbols represent the range of responses across the full gradient of reduced filtering capacity or living suspension feeder biomass.

#### 3.3.2 Reduced wind at the coastal site

At the coastal site, reducing surface wind forcing by 10% while maintaining full filtration capacity led to changes in nutrient cycling and food web dynamics relative to the baseline simulation (Fig. 5a). Reduced wind forcing lowered vertical turbulence, resulting in an approximately 10% decrease in suspended sediment concentrations. The subsequently increased light availability (+7%) increased diatom production, outweighing concurrent increases in grazing by both suspension feeders and omnivorous mesozooplankton, resulting in a net increase in diatom biomass (+17%). This increased silicate uptake and led to a corresponding decline in pelagic silicate concentrations (-15%). The remaining phytoplankton groups showed mixed responses, while total primary production and chlorophyll exhibited only minor changes (<5%). Responses were more pronounced at higher trophic levels. Omnivorous and carnivorous mesozooplankton increased substantially (by ∼90% and ∼25%, respectively), although total secondary production changed little (<2%), indicating that increases reflect changes in biomass distribution and turnover rather than increased overall production. Suspension-feeder biomass increased (+31%) supported by enhanced grazing (+92%), resulting in higher reproductive output (+29%). However, increased predation by carnivorous mesozooplankton reduced suspension-feeder larvae biomass by approximately 10%. Overall, reduced wind forcing primarily influenced the system through improved light conditions, enhancing diatom production while leading to redistribution of biomass across trophic levels rather than an increase in total ecosystem production (Table 3).

**Figure 5.**
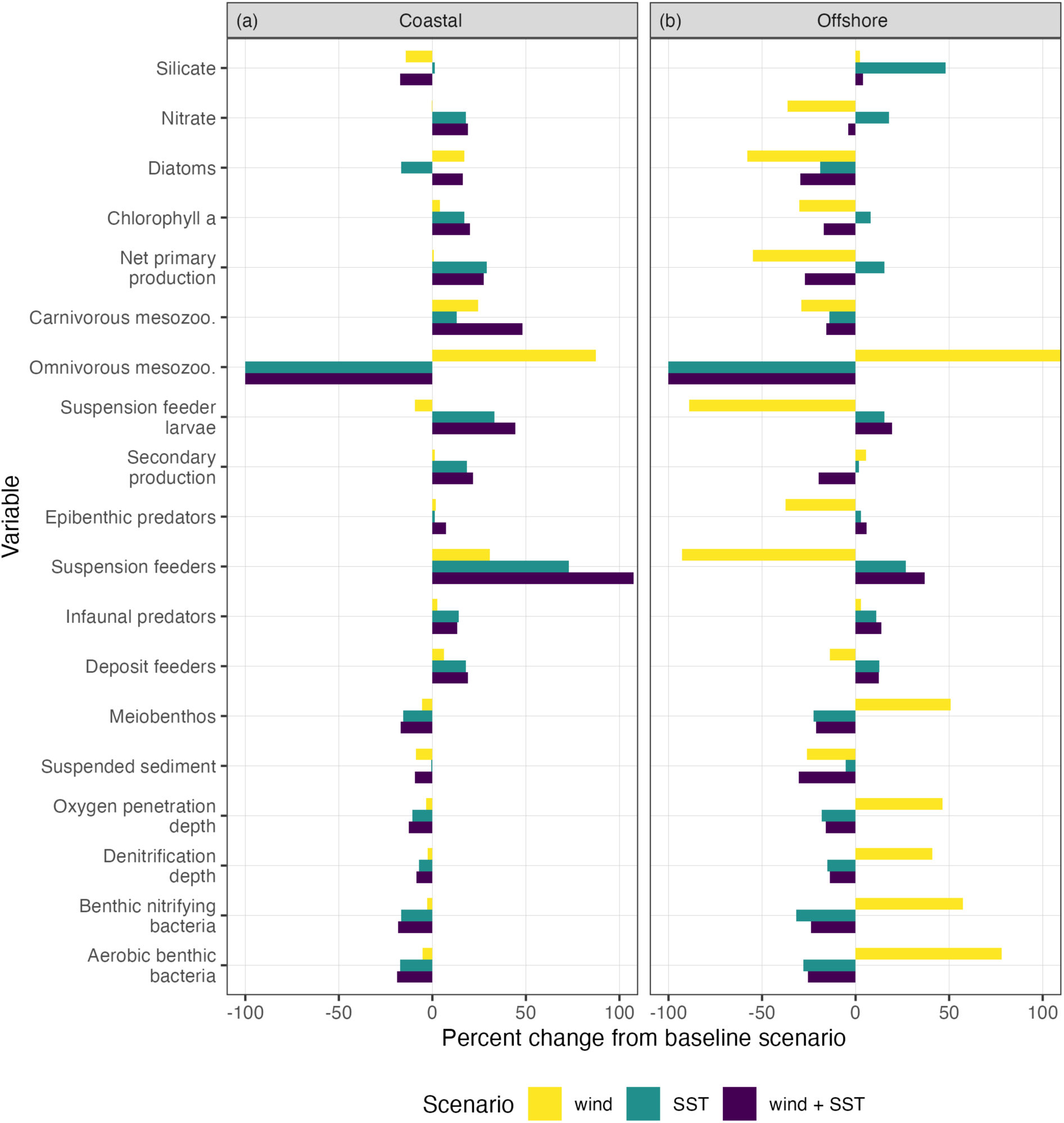
Simulated percent changes in key ecosystem variables at the coastal (a) and offshore (b) sites under reduced wind (-10%), sea surface temperature warming (+1.8 °C), and their combined effects, relative to the baseline simulation with no wind or temperature changes and full filtering capacity. Pelagic variables are from the surface layer. Bars indicate the magnitude and direction of change, highlighting differences between individual and combined drivers. Please note x axis is limited to ±100% change for visual purposes.

#### 3.3.3 Increased sea surface temperature at the coastal site

At the coastal site, increasing sea surface temperature by 1.8 °C while maintaining full filtration capacity resulted in changes in both benthic and pelagic compartments relative to the baseline simulation (Fig. 5a). In the benthos, oxygen penetration depth (-11%) and denitrification depth (-7%) declined, accompanied by reductions in aerobic bacteria (-18%), aerobic nitrifying bacteria (-18%), and meiobenthos (-16%). In contrast, most other benthic fauna increased including deposit feeders (+18%), infaunal predators (+14%), and suspension feeders (+73%) while epibenthic predators showed only minor changes (+2%). The strong increase in suspension-feeder biomass led to higher reproductive output (+61%) and increased suspension-feeder larvae in the pelagic (+33%). Increased predation by carnivorous mesozooplankton (+36%) strongly suppressed omnivorous mesozooplankton, reducing their biomass to low levels (-99%) relative to the baseline simulation. Elevated temperatures enhanced metabolic rates across functional groups, resulting in increased gross (+26%), net primary (+29%), and secondary production (+19%). Pelagic nitrate concentrations also increased by (+18%), consistent with enhanced pelagic and benthic remineralization (+25 and +13%, respectively). Overall, warming led to strong shifts in benthic community structure and trophic interactions, while also increasing ecosystem production (Table 3).

#### 3.3.4 Reduced filtration at the offshore Oyster Grounds site

At the offshore Oyster Grounds site, scenario responses were broadly similar in direction to those at the coastal site but generally occurred more rapidly and with greater magnitude. Reductions in suspension-feeding capacity propagated through the pelagic food web via similar trophic cascades as observed at the coastal site, but with amplified responses (Fig. 6a). At 50% reduced filtering capacity, suspension-feeder biomass was almost completely removed (-99%), while omnivorous mesozooplankton increased up to +300%, resulting in strong grazing pressure on diatoms (-50%) and a substantial increase in pelagic silicate (+40%). Compared to the coastal site, chlorophyll decreased (down to -12%) alongside a substantial decline in surface-layer net primary production (down to -30%), indicating that reduced production contributed directly to the lower phytoplankton biomass. In contrast to the relatively stable secondary production at the coastal site (<5% change), secondary production increased by 20% at the offshore site, consistent with the greater increase in mesozooplankton biomass. Benthic responses also diverged, with offshore increases in oxygen penetration and denitrification depths, benthic bacteria, and meiofauna, alongside declines in deposit feeders.

**Figure 6.**
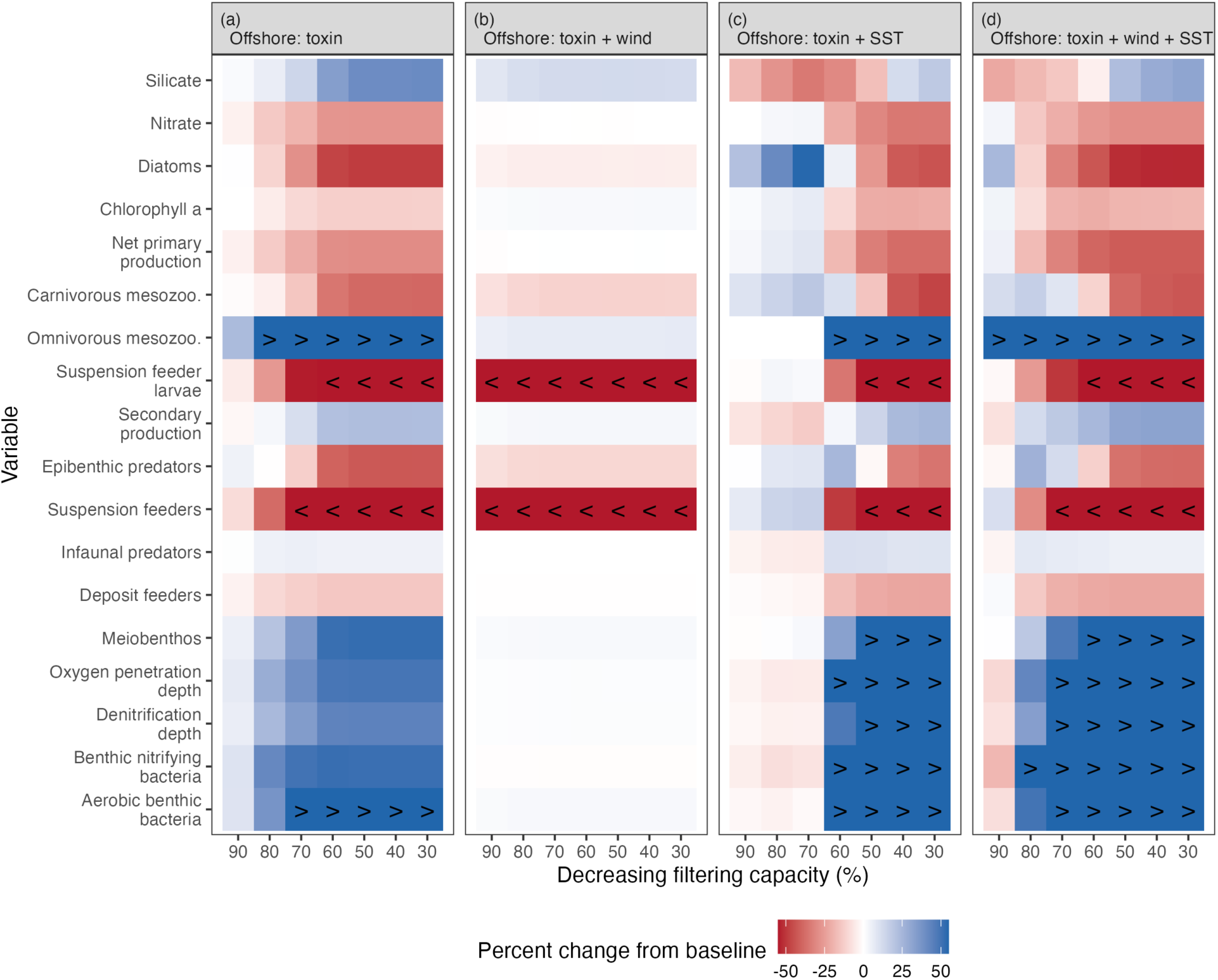
Simulated ecosystem responses to reduced filtering capacity across scenarios and their interactions at the offshore Oyster Grounds site, expressed as percent change relative to each scenario-specific baseline simulation with full filtering capacity. Pelagic variables are from the surface layer. Symbols (< and >) represent percentage changes below -50% and above +50%, respectively. Columns represent decreasing filtering capacity (from left to right) and rows represent ecosystem variables. Variable units are provided in Table 3.

#### 3.3.5 Reduced wind at the offshore Oyster Grounds site

Reduced wind forcing led to greater divergence from the coastal site (Fig. 5b). Suspension feeder and larval biomass declined by over 90%, accompanied by increased benthic activity, including greater oxygen penetration and denitrification depths and increased benthic bacteria and meiofauna. In addition, epibenthic predators (-37%) and deposit feeders (-13%) declined, a pattern not simulated at the coastal site. Responses in the water column differed markedly between the surface and bottom layers (Figs. 5b, S1). Nitrate declined similarly in both layers (-36%), whereas silicate changed by less than 5%. Despite these comparable nutrient responses, the phytoplankton community composition diverged between the layers. Diatoms declined more strongly at the surface (-58%) than at the bottom (-20%), while dinoflagellates increased substantially (>100%) in both layers. Most other phytoplankton groups generally declined at the surface but increased near the bottom, indicating a vertical restructuring of the phytoplankton community. Chlorophyll consequently declined almost twice as strongly at the surface (-30%) than at the bottom (-18%). These contrasting responses were particularly pronounced for net primary production, which declined at the surface (-55%) but only modestly near the bottom (-5%).

The zooplankton community responses also shifted, with omnivorous mesozooplankton increasing strongly in both layers (>250%), while carnivorous mesozooplankton declined more strongly at the surface (-29%) than at the bottom (-11%). Grazing by carnivorous mesozooplankton on omnivorous mesozooplankton declined (-22%), coinciding with substantially increased grazing on diatoms. At the surface, this increased grazing occurred despite little change in diatom-specific production, whereas near-bottom diatoms also experienced a substantial reduction in specific production (-50%). Secondary production increased modestly at the surface (5%) but substantially near the bottom (+40%). Overall, the reduced wind forcing produced contrasting results between the surface and bottom layers, particularly in the phytoplankton community, chlorophyll concentrations, and secondary production.

#### 3.3.6 Increased sea surface temperature at the offshore Oyster Grounds site

Responses to increased sea surface temperature followed similar patterns to those simulated at the coastal site, but were generally weaker in the pelagic and stronger in the benthic compartment (Fig. 5c). However, the increase in suspension-feeder biomass was lower offshore (approximately one-third smaller than at the coastal site), whereas nutrient responses were more pronounced offshore, with silicate increasing at the surface (+50%) and near the bottom (+75%). This silicate accumulation coincided with declining diatom biomass in both layers (Figs. 5c, S1), driven primarily by increased mortality rather than grazing, and an overall reduction (-35%) in diatom production, resulting in reduced silicate uptake. Overall net primary production showed contrasting responses between layers, increasing at the surface (+15%) but declining near the bottom (-33%). Thermal stratification increased under both scenarios (+23% under reduced wind; +11% under increased SST), while mixed layer depth changed only modestly (<4%), indicating that changes to stratification were driven by stronger vertical temperature gradients rather than changes in mixed layer depth. This increased stratification was accompanied by contrasting ecosystem responses between the surface and bottom layers, particularly under reduced wind forcing. Overall, compared to the coastal site, the offshore system was more sensitive to reduced filtering capacity and wind forcing, whereas responses to warming were generally weaker (Table 3).

#### 3.3.7 Cross-site comparison of chlorophyll phenology across individual scenarios

At the coastal site, scenario effects were primarily expressed through changes in the magnitude of the peak chlorophyll concentration rather than shifts in timing when compared to the baseline scenario (Table 4). Projected warming exerted the strongest influence, increasing the spring bloom (+23%) and nearly doubling the secondary, late-season bloom (+84%). The reduced wind scenario produced minimal changes to the spring bloom but increased the late-season peak (+24%). The increased late-season peak was associated with increased remineralization (+12%) and reduced suspended sediment (-8%), leading to improved light conditions (light extinction coefficient: -8%) when nutrients became available again later in the year. Reductions in suspension feeder filtration increased peak chlorophyll concentrations (up to +19%) in both seasons, with no clear linear relationship between the degree of filtration reduction and bloom response. Bloom timing remained relatively stable across the scenarios: peak spring blooms shifted by less 6 days, with the projected warming scenario exhibiting lower interannual variability (lower SD; Table S1). Late-season peaks advanced by approximately 2-6 days earlier, although variability in late-season peak timing was higher under some reduced filtration scenarios.

**Table 4.** Scenario-driven changes in surface-layer peak chlorophyll concentrations and bloom timing between the spring and late season at the coastal (CS) and offshore (OS) sites. The number of (+) and (–) symbols indicates the direction and relative magnitude of change. Peak chlorophyll concentrations are expressed as percentage changes (+/– = 10-49.9%; + +/– – = 50-100%; + + +/– – – = >100%), while changes of less than ±10% are denoted with (O). Bloom timing shift is expressed in days (<3 days denoted with O; +/– = 3-7 days; + +/– – = >7 days). Changes are relative to the baseline scenario, except for the Toxin + scenarios, which are relative to the specific scenarios(s) without any filtering limitation. For the toxin scenarios, symbols represent the range of responses across the full gradient of reduced filtering capacity or living suspension feeder biomass.

|  | Units | Site | Season | Wind | SST | Wind<br>+<br>SST | Toxin | Toxin<br>+<br>Wind | Toxin<br>+ SST | Toxin<br>+<br>Wind<br>+<br>SST |
| --- | --- | --- | --- | --- | --- | --- | --- | --- | --- | --- |
| Peak<br>chlorophyll<br>concen. | mg Chl m<sup>-3</sup> | CS | Spring | O | + | + | O to + | O to + | O | O |
|  |  |  | Late | + | ++ | ++ | O to + | - to O | O | O to + |
|  |  | OS | Spring | -- | + | + | - to O | O | - to + | - to O |
|  |  |  | Late | + | + | + | O | O | - to O | O |
| Bloom<br>timing | Day of<br>Year | CS | Spring | O | O | O | + | O to + | O | O |
|  |  |  | Late | O | - | - | O to + | O to ++ | O to + | O to + |
|  |  | OS | Spring | -- | - | -- | O | O | O | O |
|  |  |  | Late | ++ | O | ++ | + | O | O to + | O |

In contrast to the coastal site, both the timing and magnitude of peak chlorophyll concentrations changed at the offshore site upon comparison to the baseline scenario (Table 4). As the suspension feeder filtration was progressively reduced, the spring bloom magnitude declined in a near-monotonic manner, with peak chlorophyll concentrations decreasing (down to -22%). Conversely, reduced filtration increased late-season peak chlorophyll concentrations (+9%). Bloom timing remained relatively stable in both periods, with peak dates advancing by less than 2 days across filtration scenarios. Under the reduced wind scenario, the spring bloom magnitude declined (-39%) with decreased interannual variability (lower SD; Table S1), and the bloom advanced by nearly 11 days earlier. Projected warming also advanced the spring bloom timing by 5 days earlier but instead increased the magnitude (+29%). The late-season bloom increased under the reduced wind (+22%) and projected warming scenarios (+50%), respectively; however, only the reduced wind scenario showed a delayed late-season peak of nearly 7 days later.

Overall, the two sites exhibited contrasting responses to the individual scenarios, differing in both the magnitude and timing of peak chlorophyll concentrations (Table 4). At the coastal site, responses were dominated by changes in bloom magnitude, with projected warming exerting the strongest effect. At the offshore site, both the magnitude and timing shifted, and the direction of change varied seasonally across scenarios.

### 3.4 Combined scenario responses (interactions)

The combined (interaction) scenarios revealed both additive and non-additive ecosystem responses. Reduced filtration generally led to strong negative effects on suspension feeders, reflecting the direct effect of reduced filtration capacity, while broader ecosystem responses depended on the site-specific interactions with reduced wind forcing and increased sea surface temperature.

At the coastal site, the effects of reduced wind forcing and increased sea surface temperature were largely additive, with the temperature effect dominating (Fig. 5a). In contrast, at the seasonally stratified site, interactions between the two drivers produced more complex, non-additive responses (Fig. 5b). For example, reduced wind alone resulted in a strong increase in omnivorous mesozooplankton and associated grazing, whereas this group was nearly absent under both the warming alone and combined scenarios, indicating that the warming response dominated when both scenarios were imposed. Nutrient responses were similarly non-additive, with the offshore silicate accumulation under warming substantially reduced when combined with reduced wind. In contrast to these non-additive pelagic responses, benthic responses under the combined scenario were largely driven by warming.

At the coastal site, the combined effects of reduced filtration and wind forcing resulted in generally amplified negative responses across key ecosystem components (e.g., diatoms, carnivorous mesozooplankton, and suspension feeder larvae), indicating limited buffering and increased system sensitivity to reduced filtration (Fig. 4b). In contrast, these broader negative responses were largely absent at the offshore site. Instead, the impacts were primarily restricted to declines in suspension feeders and their larvae associated with reduced filtration capacity, although the sensitivity to this reduction was substantially higher (Figs. 6b, S2b; Table 3).

The interaction between reduced filtration and increased sea surface temperature showed generally weaker responses across variables than to reduced filtration alone, suggesting that warming partially buffered ecosystem-level impacts at the coastal site (Fig. 4c). At the offshore site, this simulated buffering effect occurred over a narrower range of reduced filtration (≥70% capacity; Figs. 6c, S2c), beyond which responses resembled those of the reduced filtration only scenario (Fig. 6a).

This buffering effect from increased sea surface temperature was also evident in the three-way interaction (reduced filtration, reduced wind forcing, and increased temperature). At the coastal site, warming continued to offset broader ecosystem responses despite persistent declines in suspension feeders (Fig. 4d). In contrast, buffering was less pronounced at the offshore site (Figs. 6d, S2d), where responses more closely resembled those reduced filtering capacity alone (Fig. 6c). Overall, the interactive effects revealed site-specific differences, with the coastal site exhibiting generally smaller overall changes than the offshore site, where responses were more pronounced and varied in direction between pelagic and benthic compartments (Table 3).

#### 3.4.1 Cross-site comparison of chlorophyll phenology across combined scenarios

Projected warming continued to exert the strongest influence at the coastal site, even under combined reductions in wind forcing and filtration capacity. Compared to increased SST alone, the addition of reduced wind had marginal impacts on the overall dynamics (Table 4). Reduced filtration in combination with the other scenarios did not produce substantial changes until filtration capacity declined to 40%. At these low filtration levels, the reduced filtration and wind forcing scenario produced dynamics in the opposite direction to the wind-only scenario (Sect. 3.3.5): spring peak chlorophyll increased (+26%), late-season concentrations decreased (-13%), and shifts in peak timing remained under 2.5 days. Adding reduced filtration to the projected warming scenario yielded changes of less than 10% in peak chlorophyll concentration relative to the warming scenario alone, with timing shifts confined to fewer than 4 days in both seasons. Compared to the scenario with only reduced wind forcing and projected warming, the three-way interaction including reduced filtration resulted in minimal spring-season changes, while the late season exhibited increases in peak chlorophyll (up to +19%) and a 4-day earlier peak.

At the offshore site, the projected warming scenario increased peak chlorophyll concentrations and shifted the bloom timing earlier; however, these effects were largely counteracted when wind and filtration capacity were jointly reduced. Wind reduction alone caused dramatic reductions to suspension feeder biomass (Fig. 5b; Sect. 3.3.4). Therefore, the added reduced filtration scenario simply compounded these existing reductions. The combined projected warming and wind reduction scenarios increased peak chlorophyll concentrations, although to a lesser extent than warming alone, and delayed both seasonal peaks by two weeks. Adding reduced filtration to the warming scenario produced nonlinear responses in the spring season: peak chlorophyll concentrations increased progressively (by up to +13%) as filtration capacity was reduced to 70%, then concentrations progressively decreased (to -36%) with further reductions in filtration capacity. In contrast, late-season peak chlorophyll concentrations decreased monotonically (to -14%). Overall, the buffering effect of projected warming was partially overcome by the additional scenarios. This was evident in the three-way interaction, where late-season changes were minimal, while spring-season responses showed nonlinear patterns consistent with the combined reduced filtration and warming scenario. However, the increases in chlorophyll concentrations were restricted to the 90% filtration capacity before concentrations progressively declined (to -34%).

Overall, projected warming exerted a strong influence at both sites, though its combined scenario effects were more pronounced at the coastal site (Table 4). At the offshore site, the influence of projected warming was partially counteracted by joint reductions in wind forcing and filtration capacity. Reduced wind had a much stronger influence at the offshore site than at the coastal site. At both sites, the addition of reduced filtration generally had less influence than the other combined scenarios.

## 4 Discussion

Offshore wind farms (OWFs) are rapidly expanding to meet growing demands for renewable energy (Díaz and Soares, 2020). As suitable shallow coastal sites become increasingly occupied, future OWF development is expected to extend further offshore into deeper coastal and shelf-sea environments (Renner, 2025). This expansion requires a robust understanding of the long-term ecological consequences of OWFs and how these may interact with ongoing climate change (Isaksson et al., 2025). Process-based ecosystem models provide a powerful framework for disentangling the mechanisms underlying ecosystem responses to OWF-related changes from those driven by climate change (Griffiths et al., 2017). Here, we examined the combined effects of OWF-related scenarios (reduced benthic suspension-feeding activity and reduced wind forcing) and climate warming (increased sea surface temperature) at contrasting coastal and offshore sites in the North Sea. Our results demonstrate that the ecological impacts of OWFs are strongly dependent on local environmental conditions, suggesting that ecosystem responses may differ substantially as wind farm development expands into deeper offshore environments.

The coastal site exhibited modest responses to the projected scenarios, largely because of its physical and biological characteristics. The comparatively lower benthic suspension-feeder biomass at the sandy coastal site relative to the muddy-sand offshore site likely contributes to the weaker benthic-pelagic coupling and more modest ecosystem responses to reduced suspension-feeder filtration in our simulations (Fig. 4a vs Fig. 6a; Tables 3, 4; Heip et al., 1992; Breine et al., 2018). The coastal site is characterized by elevated background nutrient concentrations because of its proximity to riverine inputs. Because the one-dimensional model framework does not resolve horizontal transport or lateral nutrient exchange, these simulated coastal nutrient dynamics are necessarily simplified, which may have contributed to the poorer representation of phosphate dynamics (Fig. 2; Table 1). The coastal site is shallow, with strong tidal currents, resulting in a generally well-mixed water column with little or no seasonal stratification (Ivanov et al., 2020; Ivanov et al., 2021). Because the site is already well mixed, the primary effect of reduced wind forcing was lower sediment suspension (Fig. 5a), which improved water clarity and increased light availability for phytoplankton (Van Der Molen et al., 2014; Daewel et al., 2022; Zhao et al., 2019). Our simulations do not account for drag-induced mixing around monopile foundations, which can locally offset (or exceed) reductions in surface-driven mixing (Maar et al., 2026). However, this process is likely to be of less importance at the coastal site given its already well-mixed hydrodynamic regime (Daewel et al., 2022). In contrast, reduced surface wind forcing produced much stronger ecological responses at the offshore site, where seasonal stratification plays a fundamental role in regulating ecosystem dynamics (Ruardij et al., 1997).

The offshore Oyster Grounds site responded much more strongly to the projected scenarios. The site is relatively deep (45m), characterized by muddy-sand sediments, lower background nutrients, and predictable summer thermal stratification (Van Leeuwen et al., 2015). During stratification, nutrients are progressively depleted from the warm, well-lit surface layer through primary production, increasing the importance of vertical nutrient transport for sustaining pelagic production (Ruardij et al., 1997). Reduced wind forcing and warming further strengthened thermal stratification (Sects. 3.3.5, 3.3.6), consistent with reduced vertical mixing under weaker wind forcing (Daewel et al., 2022; Maar et al., 2026). The pronounced differences between surface and bottom plankton communities suggest that this enhanced stratification altered not only nutrient cycling but also the vertical structure of trophic interactions (Figs. 5b, S1). Together with the pronounced responses to reduced suspension-feeder filtration (Fig. 6a) and scenario-driven shifts in phytoplankton bloom timing and magnitude (Tables 3, 4), these ecosystem-wide responses highlight the sensitivity of the offshore ecosystem and the strong benthic-pelagic coupling characteristic of the Oyster Grounds.

Our simulations focused on benthic suspension feeders and did not explicitly represent suspension feeders colonizing OWF structures. Although this omits an additional filtration pathway from the model, benthic suspension feeders remain a key functional group regulating benthic-pelagic coupling and nutrient cycling (Graf, 1992; Maar et al., 2007). This is especially relevant considering their susceptibility to contaminant exposure through filtration of the water column, uptake of contaminant-bound suspended particles, and exposure to sediment-associated contaminants following deposition and resuspension (e.g., Mao et al., 2011; Kádár et al., 2001; Wang et al., 2023). Incorporating turbine-associated suspension feeders could modify the ecosystem responses projected here in several ways. By intercepting phytoplankton and suspended organic matter within the water column, these communities may reduce the direct flux of pelagic food reaching benthic suspension feeders, while their biodeposition simultaneously redirects part of this material to the seabed (e.g., Slavik et al., 2019; Maar et al., 2009; Degraer et al., 2020). This redistribution of organic matter may be particularly important at seasonally stratified sites, where benthic communities rely strongly on the vertical flux of pelagic production (Van Der Molen et al., 2013; Ruardij et al., 1997). The net effect of turbine-associated suspension feeders on the system’s total filtration capacity remains uncertain, as their additional filtration in the water column may be partly offset by changes in food availability and biomass of benthic suspension feeders. If contaminant exposure subsequently reduces filtration by both turbine-associated and benthic suspension feeders, the resulting decline in total filtration capacity could leave more phytoplankton and suspended organic matter in the water column (De Borger et al., 2025), potentially amplifying the pelagic responses to reduced filtration projected here (Tables 3, 4). At the same time, reduced filtration by turbine-associated communities would decrease local biodeposition, altering the quantity and form of organic matter transferred to the seabed and potentially modifying benthic production and nutrient cycling (De Borger et al., 2021). Future model development incorporating both filtration pathways would enable assessment of how multiple filtration pathways interact to buffer, amplify, or otherwise modify ecosystem responses to contaminant exposure and future environmental change.

Our scenario projections were broadly consistent with previous modelling studies, while also highlighting differences in specific ecosystem responses. Climate-driven warming elevated metabolic and biogeochemical rates, resulting in higher primary and secondary production at the coastal site and increased primary production in the surface layer at the offshore site (Table 3). Similar responses were reported at the offshore site by Van Der Molen et al. (2013) using an earlier version of the modelling framework employed here. However, whereas Van Der Molen et al. (2013) projected declines in benthic functional groups, our simulations projected increases in all groups except meiobenthos (Fig. 5; Table 3). This difference may reflect the contrasting climate-change scenarios employed, with Van Der Molen et al. (2013) simulating conditions 30 years further into the future. Using a three-dimensional, spatially resolved ecosystem model, Daewel et al. (2022) also projected vertically heterogenous responses in net primary production under reduced wind forcing, although the vertical pattern differed from our simulation which found net primary production declined substantially more at the surface for the offshore site. These differences may reflect variation in scenario design, model dimensionality, and ecosystem model formulation (Ford et al., 2017). Despite these differences in primary production, Daewel et al. (2022) also found increased zooplankton biomass, supporting the consistency of the broader ecosystem responses.

Suspension-feeder exposure to OWF contaminant emissions was represented through reductions in filtration activity, providing a functional proxy for contaminant exposure. Given the emerging understanding of contaminant emissions from OWF infrastructure and the diversity of compounds potentially released (Hengstmann et al., 2025), we adopted a scenario-based approach that does not require assumptions about the effects of individual contaminants. This approach is particularly useful while understanding of environmental concentrations, exposure pathways, and biological responses continue to develop (Vanavermaete et al., 2026; Afshari et al., 2026). Emerging field and laboratory studies have reported variable biological responses to OWF-associated contaminant exposure (e.g., Alter et al., 2025; Zonderman et al., 2025; Ndugwa et al., 2026), further supporting the use of an exploratory range of filtration reductions rather than a single compound-specific response. Our simulations therefore examined how different magnitudes of impaired suspension-feeder filtration propagated through the ecosystem. However, reduced filtration represents only one potential response to contaminant exposure. Other physiological effects, including increased metabolic costs associated with detoxification and cellular maintenance (Widdows et al., 1995; Sokolova and Lannig, 2008), could reduce the energy available for growth and reproduction and provide additional mechanistic pathways for future model development. The modelling framework presented here can readily incorporate these additional mechanisms and compound-specific responses as new information on the biological effects of OWF contaminant emissions becomes available.

## 5 Conclusion

Our model projections identified specific, measurable ecosystem responses that could serve as early-warning indicators of ecological change associated with OWF development and climate change. The simulated trophic cascades propagated through multiple trophic levels, yet such ecosystem-wide responses are unlikely to be directly detected through routine monitoring. Instead, the functional groups (e.g., diatoms and zooplankton) and ecosystem processes (e.g., dissolved silicate concentrations and phytoplankton bloom timing and magnitude) that responded most consistently across scenarios provide practical targets for monitoring programs and may offer early warning of broader ecosystem change. Emerging technologies, such as automated plankton imaging and scanning systems (Van Walraven et al., 2025), can complement traditional monitoring by providing higher-resolution observations of these key indicators. Combined with ecosystem modelling, these approaches offer a powerful framework for detecting, interpreting, and anticipating ecological change as OWFs expand (Skogen et al., 2024).

## Code availability

Stand-alone code for GOTM can be downloaded from https://github.com/gotm-model/code.git (last access: 4th February 2025). For installation instructions, see https://gotm.net/portfolio/software/ (last access: 4th February 2025). GOTM initially developed by Burchard et al. (1999), and further developed by a group of volunteers for over 25+ years; see https://gotm.net/publications/ (last access: 4th February 2025) for references.

## Data availability

Time-series observations are available from the Rijkswaterstaat (RWS) Monitoring Waterstaatkundige Toestand des Lands (MWTL) program (https://waterinfo-extra.rws.nl/monitoring/).

## Supplement link

Please see separate Supplemental material document.

## Author contributions

Conceptualization: BD, MP, JM. Methodology: BD, JM. Formal analysis: BD. Writing-original draft: BD. Writing-review & editing: BD, MP, JM. Funding acquisition: MP, JM. Supervision. JM.

## Disclaimer

Copernicus Publications remains neutral with regard to jurisdictional claims made in the text, published maps, institutional affiliations, or any other geographical representation in this paper. While Copernicus Publications makes every effort to include appropriate place names, the final responsibility lies with the authors. Views expressed in the text are those of the authors and do not necessarily reflect the views of the publisher.

## Acknowledgements

We thank Christos Giannopoulos (NIOZ) for providing the MWTL functional group biomass data. During the preparation of this work the author(s) used AI to assist in diversifying word choice and grammar.

