## Supplemental Material for "Projected ecosystem responses to environmental changes associated with offshore wind farms and ocean warming"

### S1. Scenario responses in the bottom layer at the offshore Oyster Grounds site

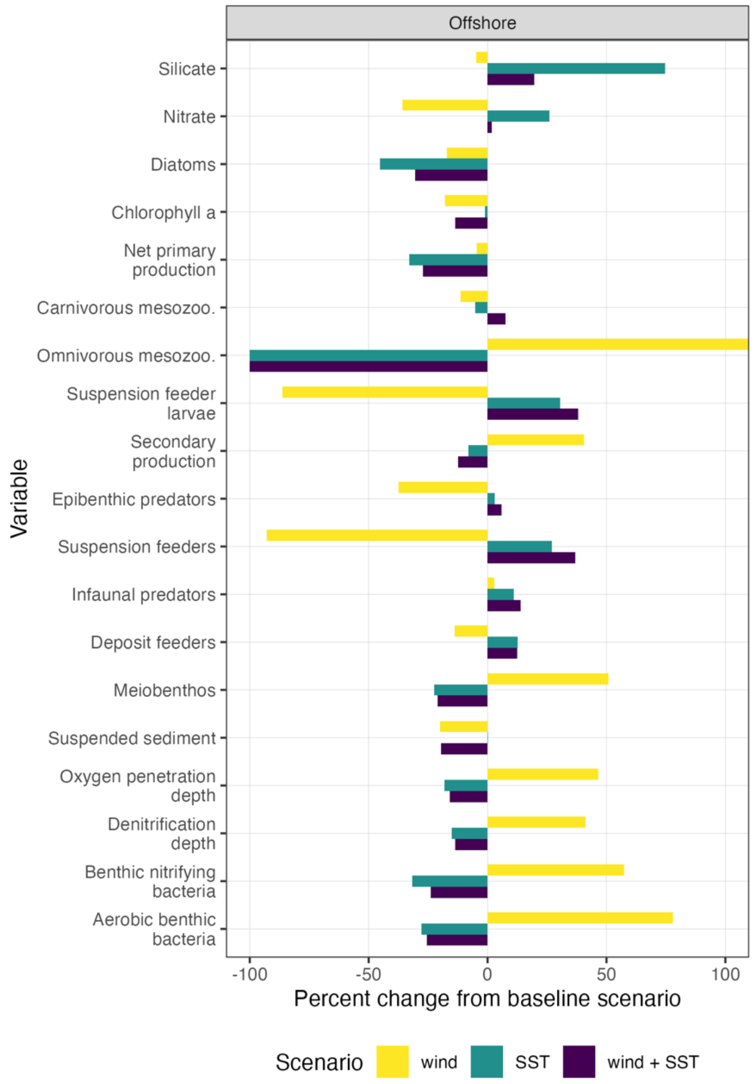


Figure S1. Simulated percent changes in key variables in the bottom layer under reduced wind (-10%), sea surface temperature warming (+1.8 °C), and their combined effects, relative to the baseline simulation with no wind or temperature changes and full filtering capacity. Please note x axis is limited to ±100% change for visual purposes.

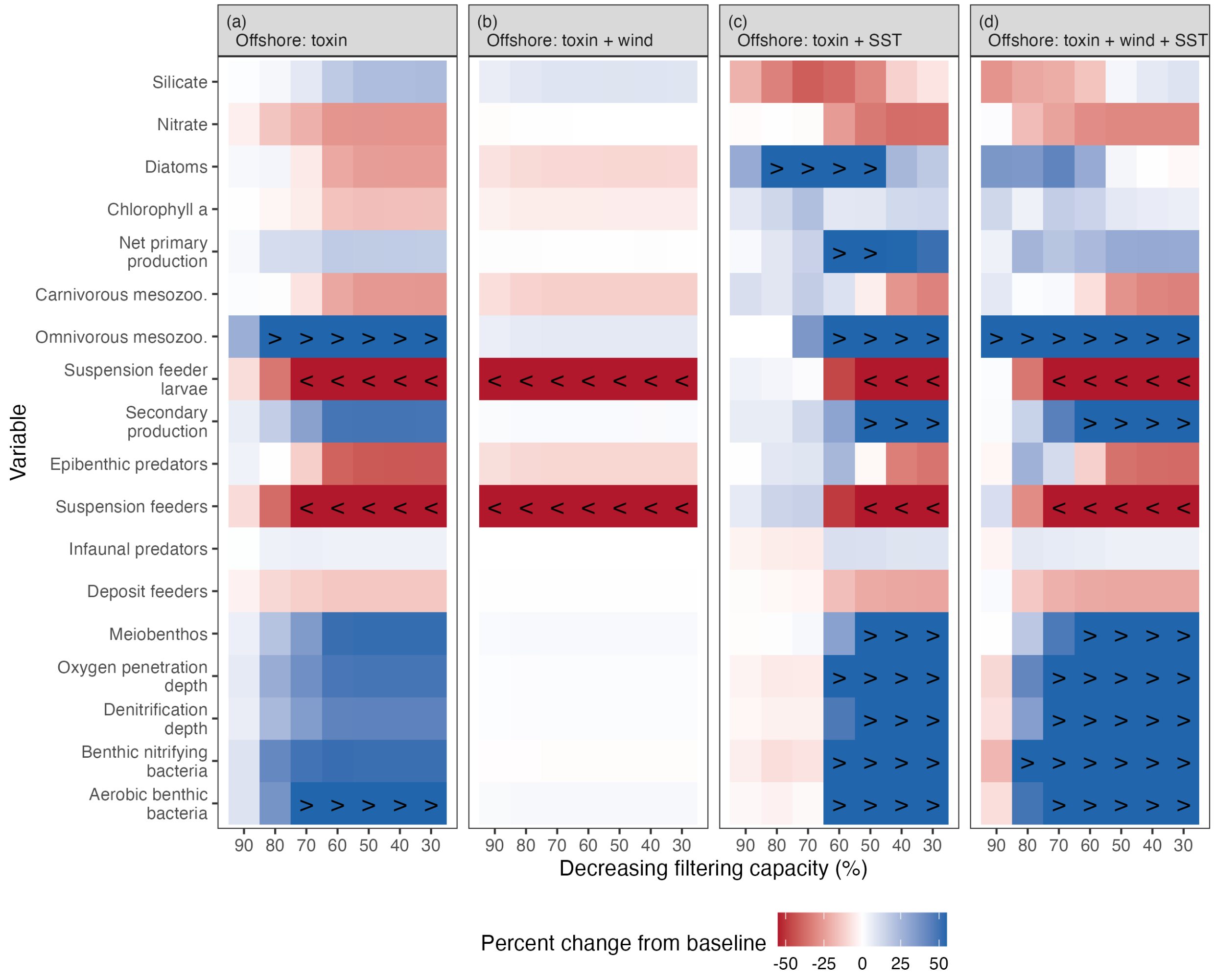


Figure S2. Simulated ecosystem responses in the bottom layer to reduced filtering capacity across scenarios and their interactions at the offshore Oyster Grounds site, expressed as percent change relative to each scenario-specific baseline simulation with full filtering capacity. Pelagic variables from bottom-layer. Symbols (< and >) represent percentage changes below -50% and above +50%, respectively. Columns represent decreasing filtering capacity (from left to right) and rows represent ecosystem variables. Variable units are provided in Table 3.

### S2. *Cross-site comparison of chlorophyll phenology across projected scenarios*

Table S1. Simulated site-specific chlorophyll phenology across the individual scenarios. The mean and standard deviation of peak bloom magnitude (mg Chl $m^{-3}$) and timing (day of year) of surface-layer chlorophyll were calculated over the 12-year simulation period (2011–2022). Seasonal differences were evaluated for the spring bloom period (1 April – 30 June) and late-season period (1 July – 31 December).

| Site | Season | Scenario | Magnitude (mean) | Magnitude (sd) | Timing (mean) | Timing (sd) |
|---|---|---|---|---|---|---|
| CS | Spring bloom | baseline | 17.9 | 4.4 | 142.4 | 16.2 |
| | | SST | 22 | 6.6 | 143.8 | 8.1 |
| | | wind | 17.4 | 4.6 | 142.6 | 13.5 |
| | | wind + SST | 21.8 | 6.5 | 142.2 | 12.8 |
| | Late season | baseline | 7.9 | 3.2 | 240.2 | 4.8 |
| | | SST | 14.5 | 6.3 | 236.8 | 4.6 |
| | | wind | 9.8 | 3.6 | 238 | 4.1 |
| | | wind + SST | 15 | 4.3 | 234.2 | 5.9 |
| OS | Spring bloom | baseline | 7.6 | 1.9 | 121.1 | 11.4 |
| | | SST | 9.8 | 2.8 | 116.7 | 9.2 |
| | | wind | 4.6 | 0.6 | 109.9 | 11.3 |
| | | wind + SST | 8.3 | 2 | 108.1 | 8.6 |
| | Late season | baseline | 1.8 | 0.5 | 304.9 | 9.7 |
| | | SST | 2.7 | 0.5 | 306.1 | 5.3 |
| | | wind | 2.2 | 0.2 | 312.5 | 5.3 |
| | | wind + SST | 2.2 | 0.5 | 318.2 | 7 |
